# Scaling up polarization-sensitive optical coherence tomography to image the whole macaque brain

**DOI:** 10.64898/2026.06.11.731509

**Authors:** MariPen Yeatts, Nirupama Chinthalapati, Bahar Baradaran, Henry M. Heinks, Emily Liao, Rachael Huxford, Vasilis Karlaftis, Taylor Grafft, Tanya K. Casta, Ayana Hellevik, Pramod K. Pisharady, Amy Howard, Jan Zimmermann, Thomas Pengo, Jonathan L. Baker, Keith P. Purpura, Matthew D. Johnson, R. Clay Reid, Franco Pestilli, Saad Jbabdi, Sarah R. Heilbronner, Taner Akkin

**Affiliations:** Department of Biomedical Engineering, University of Minnesota, Minneapolis, MN 55455; Minnesota Supercomputing Institute, University of Minnesota, Minneapolis, MN 55455; Oxford Center for Integrative Neuroimaging, FMRIB, Nuffield Department of Clinical Neuroscience, University of Oxford, Oxford, UK; Department of Psychology, Department of Neuroscience, Center for Learning and Memory, Center for Perceptual Systems, The University of Texas at Austin, Austin, TX 78712; Department of Neurosurgery, Baylor College of Medicine, Houston, TX 77030; Allen Institute for Brain Sciences, Seattle, WA, 98109; Center for Magnetic Resonance Research, University of Minnesota, Minneapolis, MN 55455; Department of Neurology, University of Minnesota, Minneapolis, MN 55455; Department of Bioengineering, Imperial College London, London, UK; Department of Neuroscience, University of Minnesota, Minneapolis, MN 55455; Brain and Mind Research Institute, Weill Cornell Medicine, New York, NY 10065

**Keywords:** axon, brain, connectivity, optical coherence tomography

## Abstract

Polarization-sensitive optical coherence tomography (PS-OCT) is a label-free imaging technique that exploits birefringence to visualize myelinated axons at micrometer resolution. However, serial PS-OCT imaging has been limited to small volumes, including tissue blocks from larger species, owing to constraints in acquisition speed, system stability, and data processing. These limitations have prevented its application to whole-brain mapping in large mammals. Here we present a scalable PS-OCT acquisition system and computational pipeline for whole-brain imaging in the rhesus macaque. The framework integrates high-throughput serial imaging with automated reconstruction and processing, enabling volumetric imaging at micrometer-scale resolution across decimeter-scale brain volumes. Using this approach, we acquired two complete macaque brains at a voxel size of 5.5 × 5.5 × 3.4 μm and an effective resolution of approximately 10 × 10 × 5.5 μm, generating multi-terabyte datasets consisting of multiple contrasts including fiber orientation information. The datasets and associated processing tools are made publicly available. This platform establishes a method for large-scale, high-resolution mapping of white matter architecture in primate brains. The resulting datasets provide a reference for validating MRI models and support the development of neurotechnological applications, including deep brain stimulation, where accurate characterization of axonal organization is required.

## Introduction

Axonal projections form the structural substrate for information transfer across functionally distinct brain regions. However, maps of these projections in larger brains, including those of humans and macaques, remain incomplete because of persistent technical limitations. Diffusion-weighted magnetic resonance imaging (dMRI) enables noninvasive whole-brain reconstruction of major axon bundles at millimeter resolution^1^, but this scale cannot resolve fine axonal architecture and is error-prone in regions with complex fiber crossings^2^. Tract-tracing methods provide high specificity and can reconstruct small groups of axons or even single trajectories^3^, but they require *in vivo* labeling, support only a limited number of injection sites, and are susceptible to histological distortion^4^. Polarization-sensitive optical coherence tomography (PS-OCT)^5,6,7^ addresses these limitations by providing whole-brain, label-free, near-distortion-free measurements at micrometer-scale resolution.

PS-OCT extends optical coherence tomography (OCT), an interferometric imaging technique that uses low-coherence light to generate depth-resolved images (**Fig. 1a**)^8,9^. By measuring changes in light polarization, PS-OCT reveals optically anisotropic tissue structures^5^. In brain tissue, myelinated axons exhibit birefringence^10^, producing measurable changes in polarization state that encode both the magnitude and orientation of local anisotropy. This contrast enables PS-OCT to distinguish highly myelinated white matter from nearly isotropic tissue^11^. It also supports tracing of axon bundles and resolution of crossing fibers within white matter with high confidence, a major limitation of dMRI (**Fig. 1b**). Unlike dMRI, which infers fiber orientations indirectly from water diffusion, PS-OCT measures signals arising directly from the myelin sheath, providing a dense anatomical reference for axonal organization^12,13^. No other current method combines these properties at comparable resolution and scale^14^.

**Figure 1:**
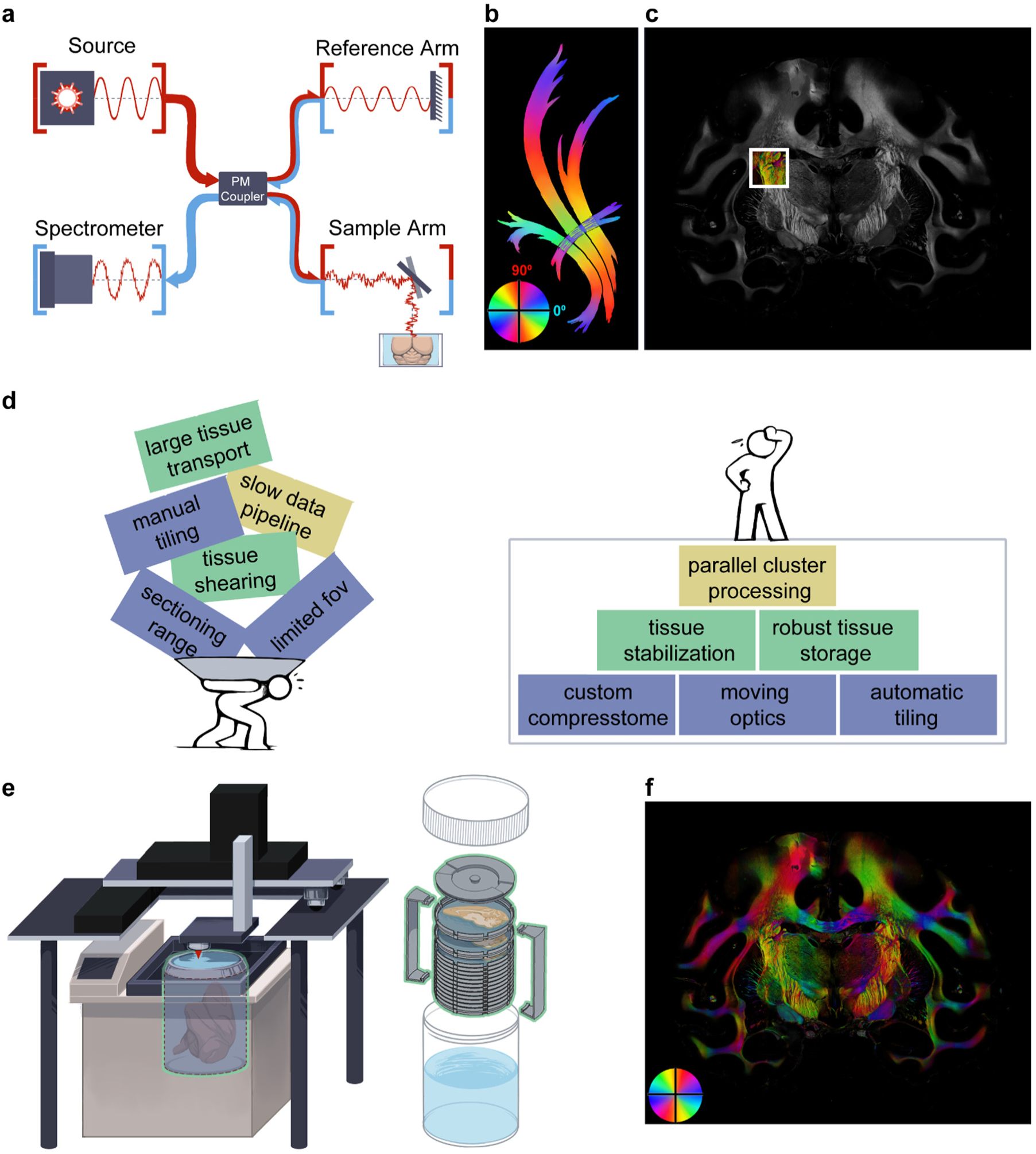
PS-OCT principles and system scaling for whole-brain imaging. **a**. Schematic of PS-OCT signal generation, showing interferometric detection of backscattered light and polarization-state changes. **b**. Illustration of fiber orientation distributions across adjacent axonal bundles including a crossing configuration. **c**. Field of view of a single tile acquisition overlaid on a macaque brain, highlighting the limited coverage of individual scans and the requirement for large-scale tiling. **d**. Engineering challenges in scaling PS-OCT to whole primate brains, including acquisition throughput, mechanical and optical stability, and data handling, with corresponding solutions implemented in the integrated system. **e**. Architecture of the scaled PS-OCT platform, including acquisition hardware and computational pipeline. **f**. Representative full coronal section of axis orientations from a complete macaque brain, demonstrating whole-brain coverage at micrometer-scale resolution.

OCT-based techniques typically achieve spatial resolutions of 1–10 μm, with imaging depths ranging from a few hundred micrometers to more than 1 mm^15^. Because OCT is label-free, it does not require tracers, exogenous contrast agents, or genetic manipulation and can be applied to both human and non-human tissue. Early studies applied OCT and PS-OCT to exposed brain surfaces, acquiring only limited fields of view^16–22^. Subsequent serial-sectioning approaches enabled volumetric imaging^12,23–30^ but were restricted to small brain regions or to small animal models. Here, we extend PS-OCT to enable volumetric imaging at the scale of the primate brain.

Previous serial PS-OCT systems were constrained by long acquisition times and physical slicing restrictions and therefore focused on rodent brains or tissue blocks from larger brains ^12,24,28,30^. Our objective was to develop an acquisition system and processing pipeline capable of whole-brain imaging in the macaque. The macaque provides a critical bridge between rodent and human neuroscience: tract-tracing is feasible, and its brain shares well-established homologies with the human brain^31^. These features make it a key model for linking microstructural organization to systems-level function and to translational applications. For example, studies of Parkinson’s disease in macaques have informed deep brain stimulation (DBS) strategies targeting the basal ganglia. Computational models further show that stimulation efficacy depends strongly on the orientation and position of electrodes relative to targeted axons^32–34^. Methods that resolve axonal organization with higher fidelity than dMRI therefore have direct translational relevance.

Here, we present a scalable PS-OCT acquisition and processing framework for whole-brain imaging in the macaque. This approach enables dense, micrometer-resolution mapping of axonal organization across the entire brain, providing a bridge between sparse tract-tracing and indirect MRI-based methods, and establishing a foundation for high-resolution, whole-brain connectomics in large brains.

## Results

We developed a toolkit for large-scale acquisition and dissemination of PS-OCT data at the level of entire macaque brains. The toolkit integrates a hardware system with a computational processing pipeline. We first outline the challenges associated with scaling PS-OCT to large primate brains and then describe the design of the acquisition system.

Whereas previous systems were optimized for rodent brains or small blocks of tissue, whole-brain imaging in the macaque requires an approximately 200-fold increase in volumetric coverage. This expansion transforms acquisition from single-section imaging to large-scale serial imaging, requiring more than 20,000 volumetric scans per brain (**Fig. 1c**). Achieving this scale introduced several constraints, including limited field of view (FOV) imposed by fixed optics, restricted sectioning range of conventional instruments, tissue deformation during sectioning of large samples, limited throughput of existing data-processing pipelines, and the need for robust containers for storage and transport of large specimens (**Fig. 1d**). Using the integrated system described below, we acquired images from two complete macaque brains while preserving tissue integrity.

To address these constraints, we developed a new PS-OCT acquisition system that increases imaging capacity. Previous implementations^12^ relied on fixed optical components positioned adjacent to a vibratome, with linear actuators used to shuttle the sample between imaging and sectioning positions. In these systems, the vibratome imposed strict limits on sample dimensions, determined by the cutting range and blade clearance, such that the sample height could not exceed the distance between the base of the chamber and the blade, nor could its lateral extent exceed the cutting range (**Supplemental Fig. 1**).

To enable whole-brain imaging, we replaced the vibratome with a compresstome (VF800, Precisionary Instruments; modified, see Supplemental Materials), one of the few sectioning systems capable of accommodating macaque brain volumes without requiring freezing or paraffin embedding, which might alter tissue properties. This change necessitated a redesign of the imaging configuration, including the transition from fixed to mobile optical components and the development of a revised tissue preparation protocol (see Methods: Sample preparation).

To transition to moving optics, a major solution was to build an imaging platform above the compresstome. To do so, we condensed the sample arm onto a 6”x6” aluminum plate (scan head, **Fig. 1e**). The platform housed three long-range, computer-controlled, motorized stages (LTS150C, Thorlabs Inc.) to move the scan head in x, y, and z directions. The optical setup for the interferometer was similar to that reported in previous studies^12^. In short, light from a superluminescent diode (center wavelength: 840 nm, bandwidth: 50 nm) was polarized and coupled into a PMF axis. A PMF-based coupler split the light onto the sample and reference arms (**Fig. 1a**) and combined the returning light. These arms included beam samplers to form additional light paths for generating a calibration signal to dynamically remove an offset from the axis orientation measurements^28^. Light mixed in the coupler yielded interference that was detected by a custom spectrometer. Light spectra corresponding to the orthogonal axes of the PMF were focused side-by-side on a line-scan camera (Octoplus, e2v), which provided raw PS-OCT data. This camera has a row of 2048 rectangular pixels (10 μm x 200 μm) contributing to the long-term stability of the spectrometer.

Custom acquisition software was written to operate the imaging system. To improve speed and consistency for large samples, automatic tiling was incorporated into the software. The movement distance between tiles was calculated from the scan size and a specified overlap percentage, generalizing tiling for any sample array. With a camera speed of 10 μs per A-line, and accounting for save time, motor movement, and tissue sectioning, we achieved a data acquisition speed of 125 tiles per hour for any given section (example of a large section with 99 tiles **Fig. 1f**). This increased throughput demanded a reliable way to verify image quality and prevent data loss. We achieved this by adding real time images of downsampled enface images to the acquisition display.

### Sample preparation and storage

Daily scanning sessions typically lasted 8 hours. To prevent tissue degradation during and after the scanning session, the system incorporated a cooling system to chill the fixed sample to 10 °C, preventing long-term degradation. The combination of cooling and agar gel embedding allowed us to pause imaging overnight while upholding tissue integrity.

The maximum sample size of a compresstome can section is controlled by the specimen tube volume. With a 120 mm diameter and 175 mm depth, our custom compresstome can accommodate a sample as large as a whole human brain hemisphere. The standard protocol to embed a sample in a compresstome is to pour agar gel into the specimen tube, over the tissue, and allow it to set. However, with larger tissue, the sample would slump out of the desired imaging position as the gel set, and was prone to voids in deep sulci. To address these issues, gel injection (**Fig. 2a**) and a mold pour were added (**Fig. 2b**).

**Figure 2:**
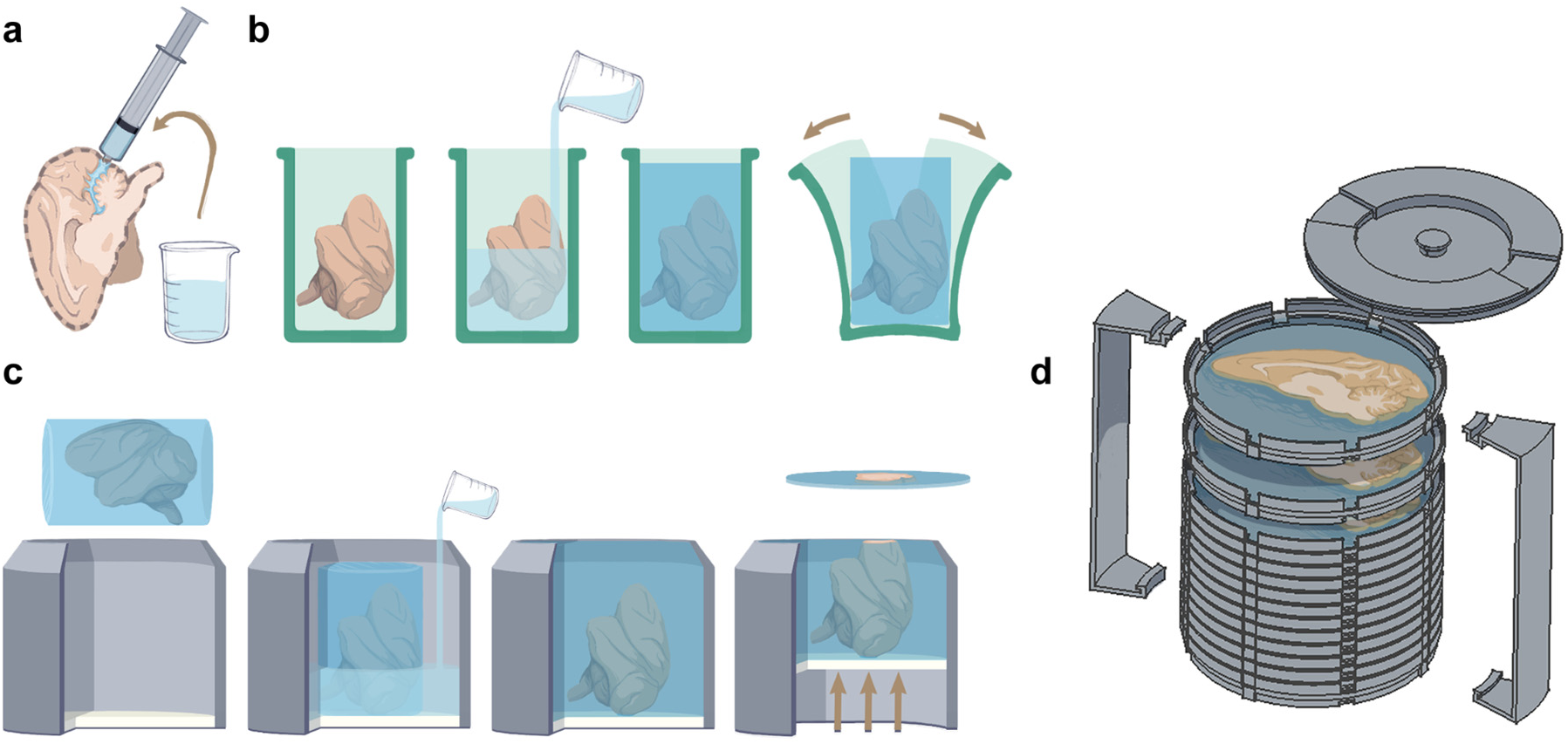
Gel embedding process. **a.** Liquid agar gel injected into spaces prone to air pockets. **b.** A whole brain positioned coronally in the silicone mold, and liquid agar gel poured to embed the tissue in the desired orientation. The gel was allowed to set before it was removed from the mold. **c.** The embedded tissue from b glued to the compresstome plunger. A second gel pour stabilized the sample in the specimen tube completing preparation for sectioning. **d.** A custom container array to store brain sections for shipment and eventual histology.

To achieve consistent tissue support during imaging and cutting, it was critical to identify a suitable agarose gel compound. We embedded the brain in agarose gel in three stages. In the first stage, a blunt, needleless syringe was used to inject approximately 100 ml of warm, liquid agarose gel into areas difficult to penetrate with submersion, including the space between the cerebellum and the occipital lobe, the central sulcus, and the interhemispheric fissure (**Fig. 2a**). The gel was allowed to briefly set, at which point stage two began (**Fig. 2b**). In this stage, the brain was placed into a silicone mold to hold it in the correct position for sectioning, and the remaining gel was poured. The mold was placed inside an incubator set to 40 °C for 60 minutes, allowing the gel to penetrate the brain. The mold was removed from the incubator and allowed to cool to room temperature prior to demolding the sample. An additional 500 ml of agarose gel was prepared for stage 3 (**Fig. 2c**) and kept at 100 °C until poured. The embedded sample was attached to the plunger of the compresstome with a cyanoacrylate glue applied to the flat surface of the gel. After allowing 5-10 minutes for the glue to cure, the additional agarose was carefully poured around the sample to fill the specimen tube, fully embedding the brain into the compresstome for sectioning and imaging. Once set, the surface of the gel was then sectioned until the tissue was visible, providing a smooth imaging surface to begin acquisition.

A section thickness of 200 µm was chosen to minimize the effect of light attenuation within sections in the depth direction and limit the total number of sections to image and store. The best sectioning settings are variable based on mechanical properties of the tissue. A speed of 0.5 mm/sec (speed setting 2) and an oscillation rate of 68 Hz (oscillation setting 8) yielded the most consistent quality, minimizing tissue deformation and maximizing section integrity. Each section was collected from the bath and transferred into a custom stackable dish filled with 10% phosphate-buffered saline (PBS) to maintain section flatness and ordinal number for future histology comparison with the PS-OCT images (**Fig. 2d**). Once complete, each stack of dishes was secured with a base, clips, and a lid and placed in a 1000 mL specimen jar (ThermoFisher catalog number 2116-1000) filled with PBS for storage until shipment (**Fig. 1e**) for subsequent study.

### Data Processing Pipeline

Each brain generated 80-115 TB of raw data. Leveraging the Minnesota Supercomputing Institute (MSI), the data was accessed directly from cloud storage and processed in parallel by section.

Data were processed in two main stages (**Fig. 3**). The pre-processing stage started by loading and reshaping the data to the appropriate tile dimensions (1000×1000) and calculating the spectral background by averaging in the x and y to one A-line (**Supplemental Fig. 2**). The background was subtracted, and the two channels were separated. After background removal, the signal was zero-padded and interpolated into linear k-space. Dispersion compensation and a hamming window were then applied. Finally, the signal was Fourier transformed into the spatial domain, resulting in two channels of complex depth profiles (CDP) per A-line, represented by *A*(*z*)*e^ie(z)^* (**Fig. 3**). The CDP was the end product of the pre-processing stage, maintaining all necessary information to derive PS-OCT contrasts such reflectivity, cross-polarization, phase retardance, and axis orientation as depicted by the equations in the processing section of **Fig. 3**.

**Figure 3:**
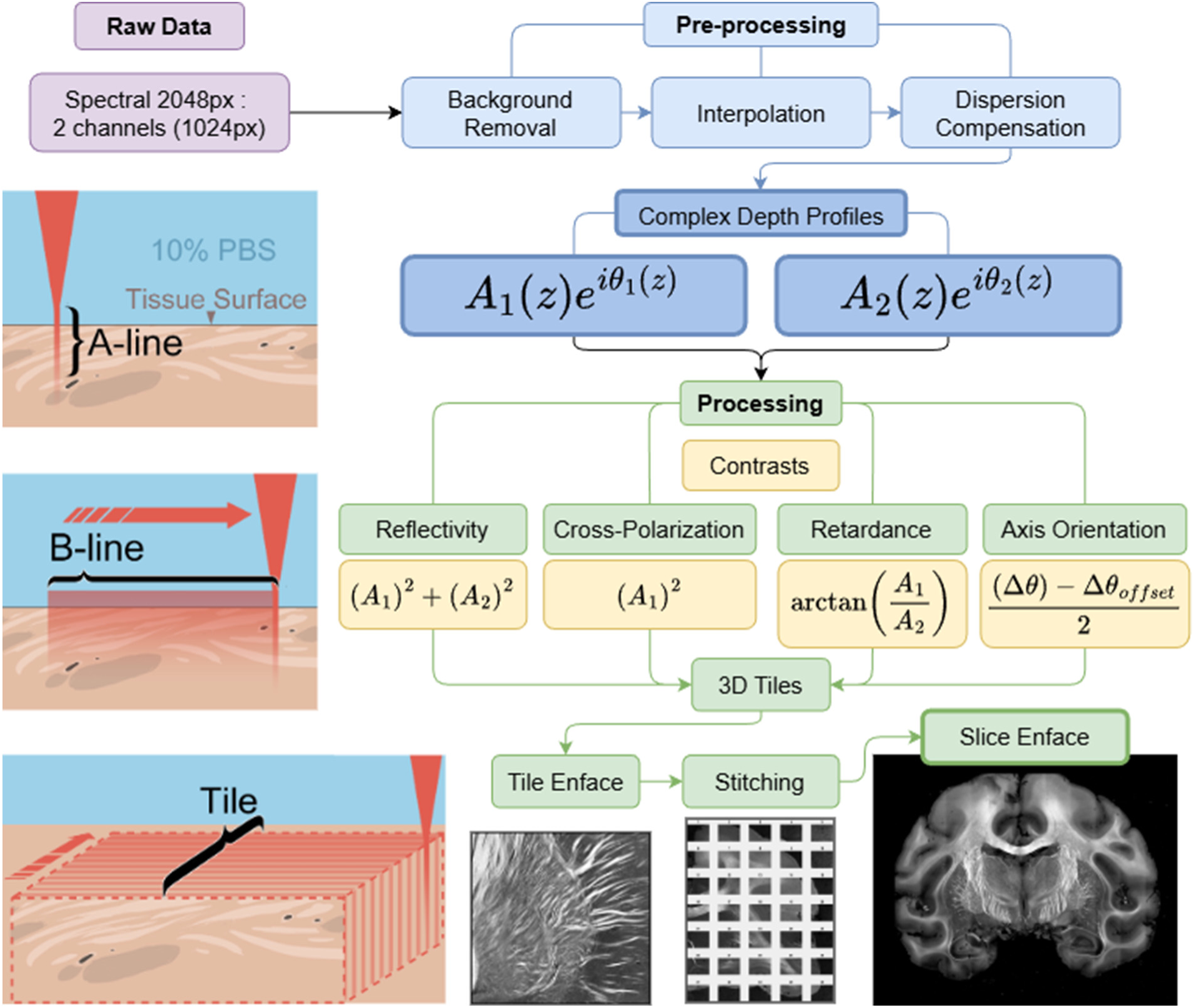
Processing diagram. Each A-line has 2 channels which correspond to 1 scanline in depth. Performing a single scan in the x direction creates the B line (1000 A-lines). A raster scan (tile) generates 1000 B-lines (or 1000 x 1000 A-lines). Pre-processing takes in raw data and converts it to complex depth profiles, translating it from the spectral to the spatial domain. Processing extracts individual contrasts from complex depth profiles.

For each brain section, we generated four contrasts: reflectivity, cross-polarization, phase retardance, and axis orientation (**Figs. 4a and 5a**). To create an enface image for each contrast, a weighted average was taken along the depth of each tile as visualized in the left of **Fig. 3**. Retardance and orientation were kept in the complex domain to avoid unwrapping errors during averaging. The tile enface images (sized 1000 x 1000 pixels) were then stitched with a 10% overlap and a simple linear blend to generate the composite slice enface images for each contrast. The resulting enface images provided an impressive level of detail with regard to brain structure.

**Figure 4:**
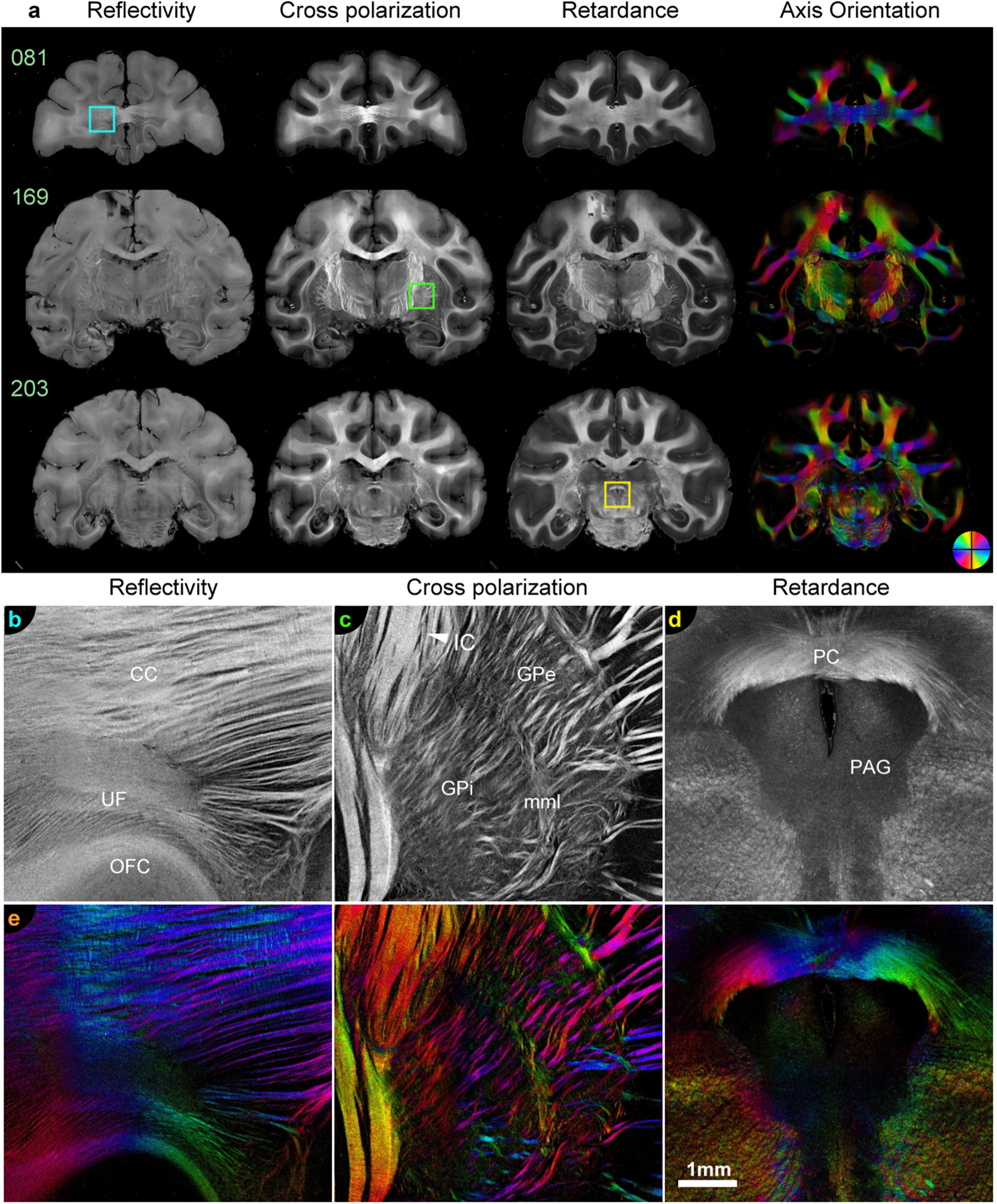
Coronal macaque brain (Subject M) enface contrasts and zoom in. **a.** Four contrasts across three slices **b.** Enlarged reflectivity image of fibers of the prefrontal cortex from slice 81 as marked by the cyan rectangle. Fibers from the prefrontal cortex cross to the contralateral hemisphere via the corpus callosum (CC); fibers of the orbitofrontal cortex (OFC) and uncinate fasciculus (UF) commingle. **c.** Enlarged cross-polarization image of the globus pallidus from slice 169 as marked by the green rectangle. The globus pallidus internus (GPi) is easily differentiable from the globus pallidus externus (GPe) and putamen. Fine details of the fibers of the internal capsule (IC), medial medullary lamina (mml), and lateral medullary lamina are also visible. **d.** Enlarged retardance image of fibers of the midbrain from slice 203 as marked by the yellow rectangle . Fibers of the posterior commissure (PC) are visible next to the periaqueductal gray (PAG). **e.** Axis orientations of the same regions of interest in b,c, and d.

**Figure 5:**
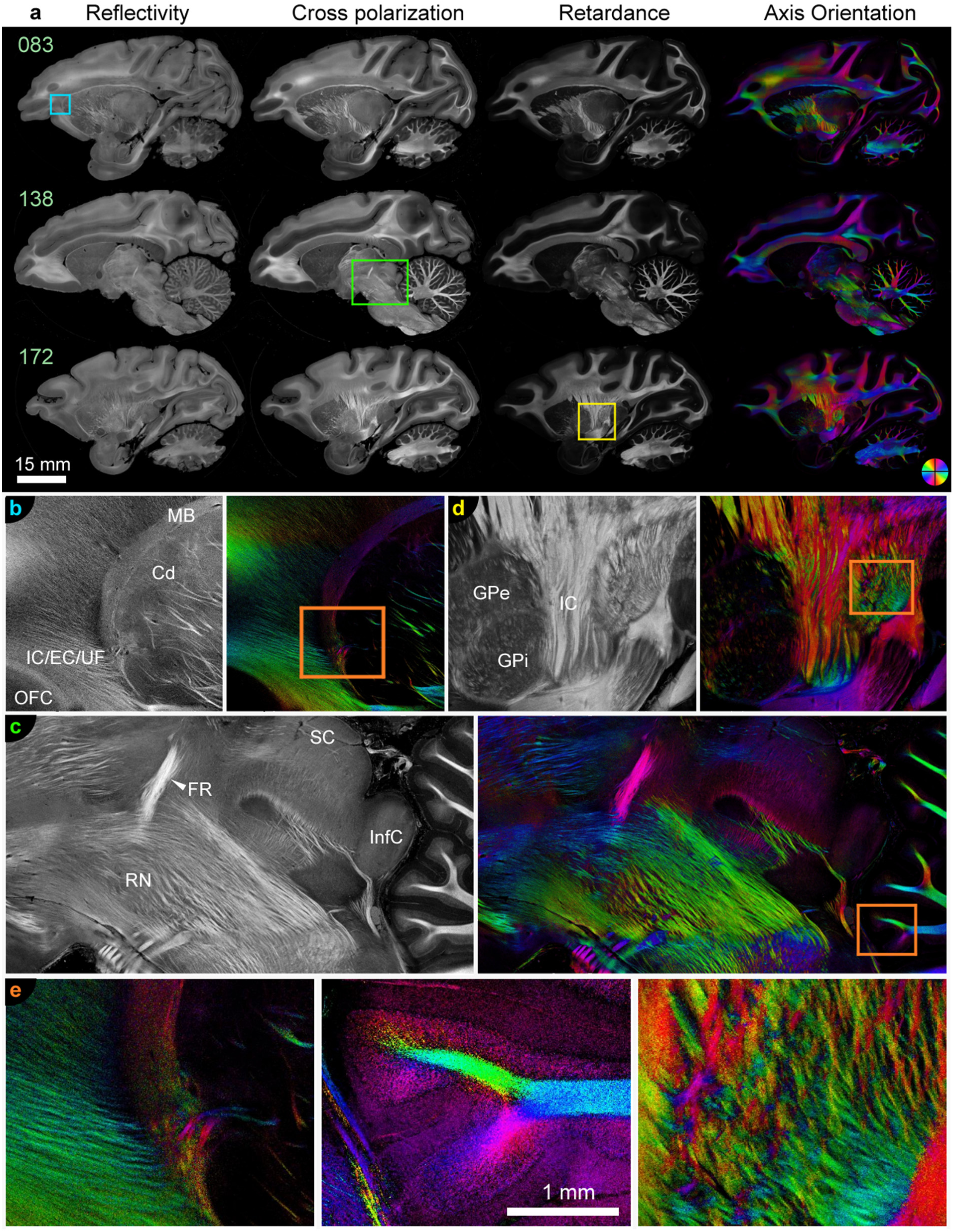
Sagittal macaque brain (Subject V) enface contrasts and zoom in. **a.** Four contrasts across three slices **b.** Reflectivity zoom in blue rectangle. Fibers from the prefrontal cortex enter the caudate nucleus (Cd) at its head. Muratoff’s bundle (MB) contains axons streaming anterior-posteriorly to reach the appropriate level of the caudate for termination. A complex mixture of bundles that, at more posterior positions, will form the internal capsule (IC), external capsule (EC), and uncinate fasciculus (UF) can be seen gathering ventral to the head of the caudate **c.** Cross polarization zoom in green rectangle. The section shows complex fibers through a sagittal section at the intersection of the diencephalon, midbrain, and pons. The superior colliculus (SC) and inferior colliculus (InfC) can be clearly segmented. The fasciculus retroflexus (FR), traveling primarily dorsal-ventrally, is bright. The complexity of the fibers of the midbrain reticular formation can be appreciated. **d.** Retardance zoom in yellow rectangle. Fibers of the IC traveling primarily dorsal-ventrally are very bright. It is possible to segment the globus pallidus internus (GPi) and externus (GPe) from each other. **e.** Axis orientations zoom in as marked by the orange rectangles in b,c,and d

Reflectivity images - sensitive to refractive index mismatches and scattering microstructure (**Figs. 4b and 5b**) - allow for segmentation of tissue from the background and visualization of tissue boundaries such as ventricles and large blood vessels. Together, cross-polarization and retardance images provide fine-grained segmentation of gray matter from white matter as well as information about structural variation, such as bundling and fasciculation, within white matter (**Figs. 4c-d and 5c-d**). There is also interesting orientation-dependent contrast. Fibers lying within the imaging plane (and therefore approximately perpendicular to the incident beam) produce stronger signal and appear hyperintense, whereas fibers oriented out of the plane, with a component parallel to the beam, appear relatively hypointense. Although PS-OCT provides sensitivity to three-dimensional fiber orientation, reliable estimation of fiber inclination remains non-trivial and is an active area of investigation. Consequently, the orientations shown here are restricted to the two-dimensional in-plane components (**Figs. 4e and 5e**).

The Supplemental Videos for Subjects M and V show the axis orientation images with the brightness controlled by retardance values. To remove noise from voids, such as in the ventricles, the retardance was masked by thresholded reflectivity.

A 3D slide deck was formed by stacking the cross-polarization slice enface images and performing co-registration of slides. To co-register the images, each slide was zero-padded to the dimensions of the largest slide and then downsampled by a factor of 10. Translations and rotations between successive PS-OCT slides were estimated using FSL’s FLIRT^35^. The estimated transformations were concatenated such that all slides were aligned to the largest, central sections. The result was a complete 3D volume (**Fig. 6 and Supplemental Fig. 3**). This 3D volume is used for co-registration with MRI data.

**Figure 6:**
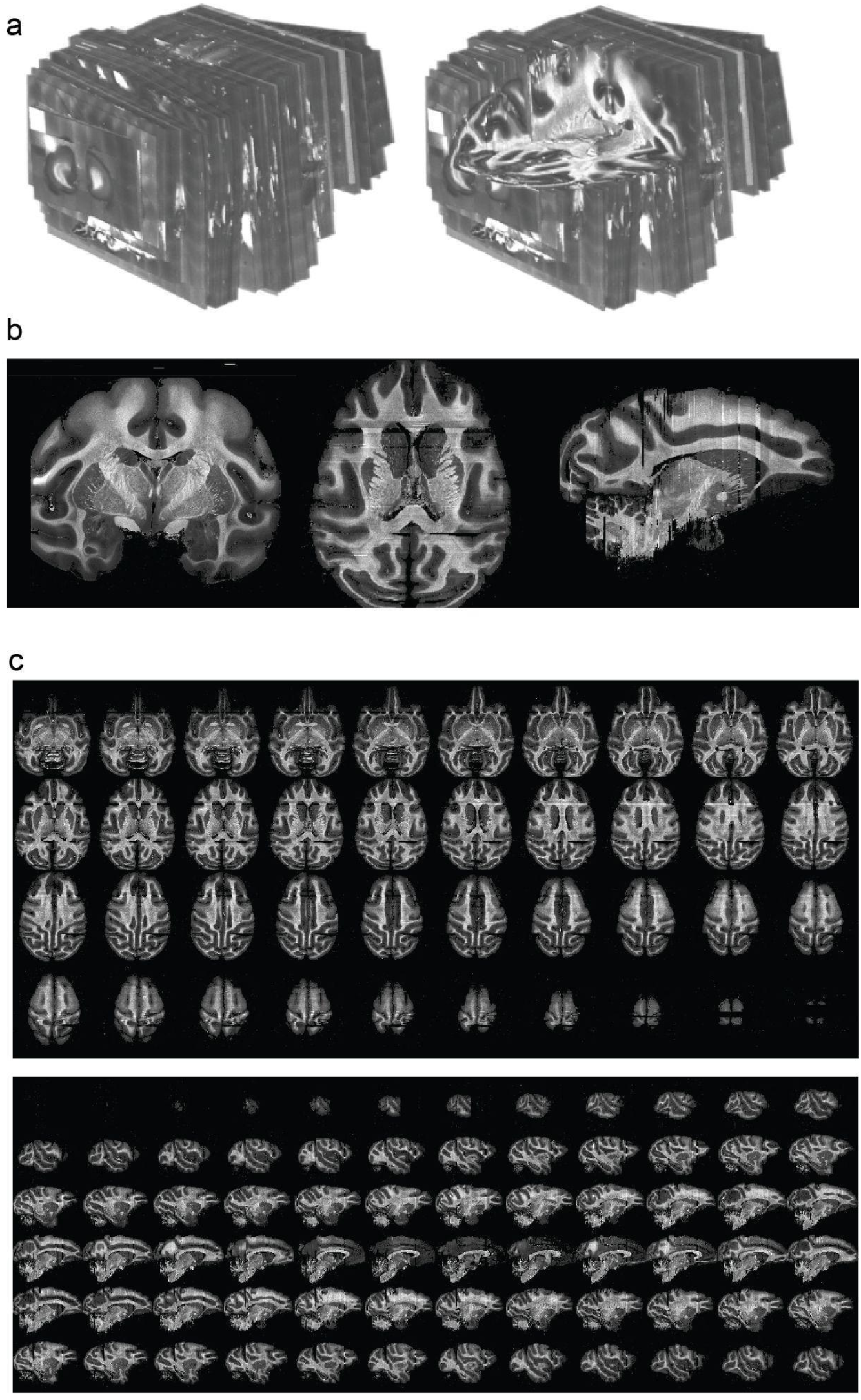
Alignment across sections produces 3D PS-OCT volumes. **a.** Slide-deck post alignment. **b**. Coronal (imaging slice) and axial views showing good between-slice alignment. **c**. Sagittal (top) and axial (bottom) views showing the good between-slice alignment of anatomy.

Data and processing code are publicly available at <u>brainlife.io.</u>

## Discussion

Here, we provide a new method of acquiring a whole macaque brain with PS-OCT and the resulting images of two complete brains. The unique space occupied by PS-OCT, as a label free imaging technique with spatial resolution two orders of magnitude finer than MRI, provides a bridge between tract-tracing and MRI. By highlighting white matter at a voxel size of 5.5 × 5.5 × 3.4 µm, we have demonstrated our data’s potential future value in connectomics and microstructural analysis. A PS-OCT system capable of imaging large brains was essential to this end.

The white matter of the primate brain occupies roughly half its volume, and the axonal connections it contains are the structural basis of long-range communication. As corroborated with tract-tracing and myelin stains, brain connectomics can be studied via white matter trajectories^14^. However, there remain major gaps in the literature in our understanding of white matter organization, particularly for large primate brains. In this study, we created a method to obtain the data needed to help fill these gaps and demonstrated its viability by imaging two complete macaque brains, one in the coronal and one in the sagittal plane.

Our next goal will be to image an entire human hemi-brain. The human brain is roughly 13 times the size of the macaque brain. While we anticipate that this will lead to additional challenges in data size, we are working to refine our data file type and structures to compensate. The completed macaque brains are currently undergoing multimodal analysis, as Subject M and Subject V have been multimodally imaged by high-field in-vivo dMRI, high-field ex-vivo dMRI ^36,37^, PS-OCT and light sheet microscopy.

A limitation of our current method is the fact that the measured orientations are in the 2D (xy) plane, missing the through plane component (polar angle or inclination angle). This has been addressed in other PS-OCT systems by introducing a second scanning angle^28,38^. However, this solution doubles the amount of data saved, significantly increases the acquisition time, and currently operates with fixed optics. We are investigating computational methods to extract the through plane angle from our current data to complete estimation of 3D orientations. One possibility is that the through plane component might be estimated from a normative brain acquired in the perpendicular plane. This is one advantage of our current dataset, in which one brain was imaged coronally and another sagittally.

We have demonstrated successful large-scale PS-OCT imaging of the macaque brain. With this system, we will continue to gather data from primate brains and interrogate the fine white matter structures in regions such as the basal ganglia, brainstem, and the prefrontal cortex. We expect that this method will continue to develop and provide a framework for large-scale imaging of additional species and other tissues.

## Methods

### Sample preparation

Experimental procedures were performed with the approval of the Institutional Animal Care and Use Committee (IACUC) at the University of Minnesota. Adult rhesus macaques (n=2 males) were subjects in this study. Prior to brain extraction, both subjects had intracranial injections of tract-tracers, the results of which are not shared here. Brains were intracardially exsanguinated and perfused with 4% paraformaldehyde (PFA). The PFA was mixed with hydrogel monomer solution (HMS), as described in Xu et al., 2021^3^, to facilitate later histology. After extraction, brains were fixed overnight, washed, and stored in 10% PBS until imaging. Because of additional dMRI scans performed between extraction and PS-OCT, a variable number of days passed between extraction and PS-OCT imaging. This had no significant impact on the resulting images. To improve slice quality, extra care was taken to remove blood vessels and dura mater prior to embedding to improve slice quality.

The amount and concentration of gel varied across subjects. For Subject V, 400 ml of 3.25% wt low-melting-point agarose gel solution was prepared by heating to 140 °C under a constant stir until clear. It was kept at a minimum of 40 °C to maintain the liquid state prior to use.

For subject M, we embedded directly in the sample tube using 2% agarose gel. Due to insufficient surface area to glue the tissue to the plunger, during imaging of subject M we observed small rotations between some sections. Additionally, there was some tissue gouging in the last third of subject M due to gel softness and degradation.

To address rotation and gel degradation, different agarose concentrations were tested. Tissue was embedded in the sagittal position using a silicone mold as shown in Figure 2b with 4% agarose gel. The flat surface of the gel was glued to the plunger. The sample was then fully embedded in the specimen tube with a second agarose gel pour. This eliminated rotational misalignment. However the increased gel stiffness caused cracking during sectioning. After testing agarose gel concentrations between 2% to 4%, 3.25% agarose gel was determined to yield the most consistent sections.

Narrow passages such as the gap between the cerebellum and the occipital lobe were notably empty or partially filled in Subject M, which lead to minor sectioning artifacts. To address such deep and narrow crevices that had difficulty filling prior to the gel setting, Subject V had agarose gel injected into these areas as shown in Figure 2a. Large hollow blood vessels also had difficulty sectioning and cast shadows around the edges of Subject M’s images. Extra care was taken to remove as much of the dura and blood vessels from Subject V prior to embedding, significantly improving image and section quality.

### Optical system description

This optical system (**Supplemental Fig. 4**) is an extension of our previous system^11,28^. The light source is a superluminescent diode (Superlum S840-B-I-20) emitting low-coherent light centered at a wavelength of 840 nm with a spectral bandwidth of 50 nm. Passing through a polarizer in a fiber bench, linearly polarized light is coupled into a polarization-maintaining fiber (PMF) based 2×2 70:30 coupler (Evanescent Optics Inc). The coupler, which has FC/APC connectors on all ends, splits the incoming light into the reference and sample arms.

In the reference arm, the linearly polarized light emerging from the PMF is first collimated by a fiber collimator and then transmitted through a beam sampler. The beam sampler directs 4% of the light to a calibration path, which consists of a quarter-wave plate with the axis at 22.5°, an achromatic lens, and a mirror. In the main path, the light continues through a linear polarizer whose transmission axis is set to 45°. Then an achromatic lens focuses light onto a partial reflector (a glass wedge). This configuration ensures that light returning from the reference arm couples back into both PMF axes with equal power.

In the sample arm, a triplet collimator (Thorlabs, TC18APC-850) collimates the linearly polarized light. A beam sampler directs 4% of the light to a calibration path that is identical to the one in the reference arm. The main path comprises an achromatic quarter-wave plate (QWP), a two-axis galvanometer-based scanner (GVS002, Thorlabs), and a scan lens (Thorlabs, LSM03-BB). A custom mounting system was designed to attach and align a mounting adaptor (GCM102, Thorlabs Inc.), scanning galvo mirrors and the scan lens. This configuration with the QWP axis at 45° illuminates the sample with circularly polarized light. An anisotropic tissue alters the polarization into an elliptical state such that when the light returns to the PMF, it has unequal power between channel axes.

Light returning from the reference and sample arms interferes in the PMF coupler, which transmits the signal to the detection arm. A home-built spectrometer in the detection arm records the interference signal. The spectrometer consists of an achromatic collimating lens, a volume phase holographic grating (1200 lines/mm, Wasatch Photonics), a Wollaston prism (Custom 6°, Karl Lambrecht Corporation), a camera lens (EMERALD 2.9/100mm, Schneider Optics Inc.), and a line scan camera (OctoPlus, Teledyne e2v). This configuration enables the spectra emerging from PMF channels to be focused side by side on the same camera.

To synchronize the galvo mirror scanning and image acquisition a DAQ card (NI PCIe 6361) controlling the galvo mirrors and a frame grabber (NI PCIe 1437) controlling the camera were connected by a RTISI cable. Raw data acquired in LabVIEW were saved to local hard drives during imaging and then uploaded to cloud storage overnight between imaging sessions.

A single capture of the camera was used to compute two complex-valued depth profiles (channel 1 and channel 2) for a single lateral location (A-line in the z-direction). To capture all locations within an approximately 5.5 x 5.5 mm^2^ FOV, the light beam was raster-scanned in the x and y directions by using a pair of galvo-mirrors. The resulting tile was saved to disk in buffers of 10 scans in x-direction (B-lines) and named to identify the slice number and tile location for later reconstruction.

### Processing

A linear blend was used to stitch individual tile enface images into a slice enface image. The overlap regions of two adjacent tiles were multiplied by opposing linear gradients from 0-1 and then summed, returning the magnitude to a factor of 1. All images were taken with a 10% overlap; equivalent to 100 pixels shared between two adjacent tiles.

The axis orientation contrast was also corrected by removing the polarization imperfections from the system across a given tile as measured in the gel area. A polynomial fit was applied to a smoothed gel tile to generate a removal matrix that was then inversely scaled by the magnitude of the axis orientation for each tile. The angular offset was translated into a complex number and then removed.

## Acknowledgements

We acknowledge funding from UM1NS132201, R01MH126923, R01NS0811108, U24NS140384, and R01NS111019. We thank the entire Center for Mesoscale Connectomics for assistance with the conceptualization and implementation of this project. We thank Amazon Web Services Open Data Sponsorship Program for supporting data storage for <u>brainlife.io</u>.

