## Supplementary Materials for "Scaling up polarization-sensitive optical coherence tomography to image the whole macaque brain"

### Supplemental Methods and Results

Further detail on the limitations of the previous imaging setup, background subtraction methodology, and optical components are provided below in the supplemental figures 1,2, and 4, respectively. Additionally, the 3D slide deck of subject V is visualized similar to Figure 6 for subject M in supplemental figure 3. The customization of the VF-800 compresstome is elaborated and alternative gel fixation methods explored to further section integrity are detailed below.

#### Compresstome Customization

Once we received the compresstome, we worked with engineers from the manufacturing company (Precisionary Instruments) to improve and test blade holder designs addressing corrosion from the PBS on the oscillating mechanism and section movement during cutting. A splash guard was added to the oscillating mechanism, and the front of the blade holder was curved and windowed. Additionally, a cooling system was implemented, with a Lytron Kodiak series recirculating chiller to regulate the sample temperature. The chiller was set to 8 °C while sample bath temperature was fluctuating around 10°C. Slice quality and sample integrity improved with these changes, which were implemented between Subject M and Subject V. Sections were collected intact from the imaging bath and placed into long-term storage as visualized in figure 2d.

#### Alternative Gelling Investigation

To maintain compatibility with downstream tissue clearing, we investigated the feasibility of hydrogel embedding using the HMS formulation without paraformaldehyde (PFA). Standard SMART protocols include PFA in the hydrogel monomer solution, where it crosslinks tissue biomolecules and anchors the polymer network within the sample[3]. However, concerns about over-fixation and its potential impact on clearing efficiency motivated efforts to reduce or eliminate PFA during gelation.

We first evaluated bulk gelation in the absence of tissue using the HMS formulation without PFA. To optimize gel formation, polymerization conditions were systematically varied, including temperature and duration. Although gelation could be achieved under extended conditions, the resulting hydrogels were mechanically weak and inconsistent. When these conditions were applied to test tissue, the hydrogel failed to adhere and did not form a stable, tissue-integrated matrix. While acrylamide polymerization likely occurred, the absence of PFA prevented effective covalent coupling between the forming hydrogel and tissue components.

These results are consistent with the underlying chemistry of CLARITY-type methods. PFA forms methylene bridge crosslinks between proteins and facilitates interactions between tissue biomolecules and acrylamide monomers, enabling formation of a stable hydrogel–tissue hybrid. In its absence, the polymer network remains unanchored, resulting in poor mechanical integrity and loss of structural support during handling[39]. Modifying polymerization conditions alone was insufficient to compensate for the lack of chemical crosslinking.

Together, these findings indicate that inclusion of PFA, or an alternative crosslinking strategy, is required for effective hydrogel embedding in large brain samples, even when downstream applications impose constraints on fixation.

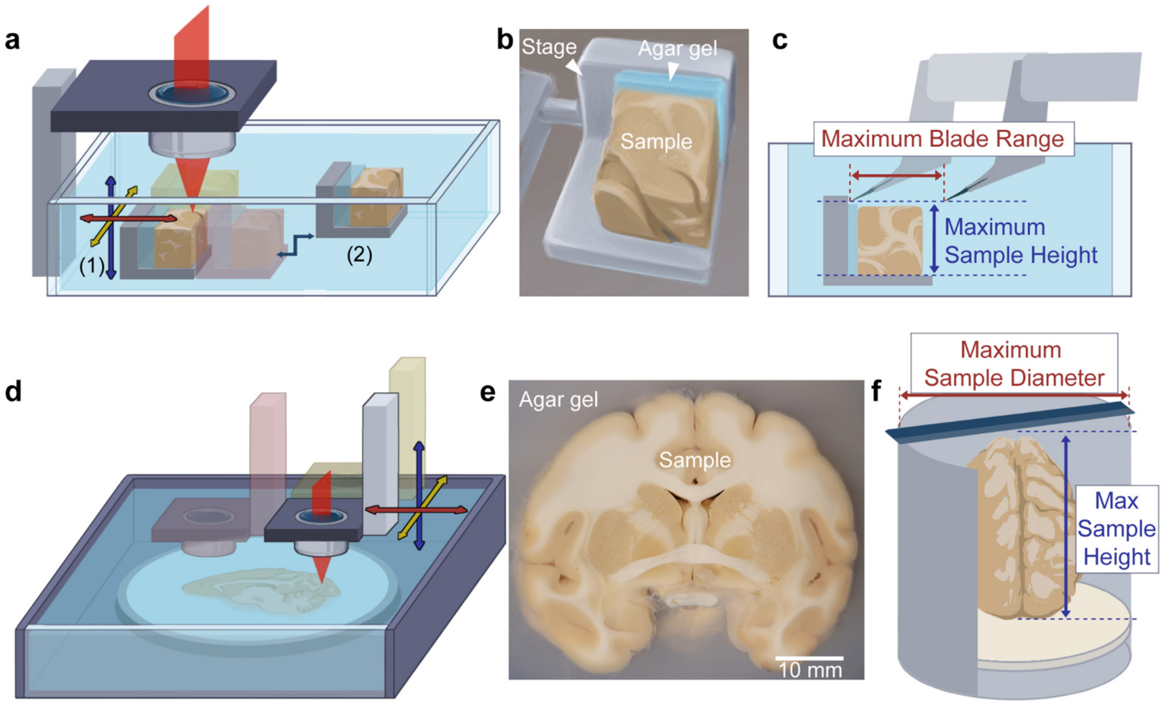


**Supplemental Figure 1: Vibratome Limitations. a.** Fixed optics and a moving sample. (1) referring to the imaging position with a maximum translation of 20 mm in the x and y (red and yellow arrows). (2) slicing position to engage with the vibratome blade **b.** Motor controlled aluminum stage. Agar gel backing to prevent blade damage from hitting the stage during slicing. Sample of a blocked section of macaque brain around the thalamus. **c.** The maximum blade range is the total cutting distance of the vibratome. The maximum sample height is determined by the distance from the stage surface, where the sample is glued, to the blade. **d.** Fixed sample and moving optics **e.** Photo of Subject M between data acquisition. **f.** The maximum sample diameter is less than the opening of the specimen tube to ensure the sample does not touch the specimen tube walls. The maximum sample height is the distance from the cutting surface (top of the specimen tube) and the plunger surface in the lowest position.

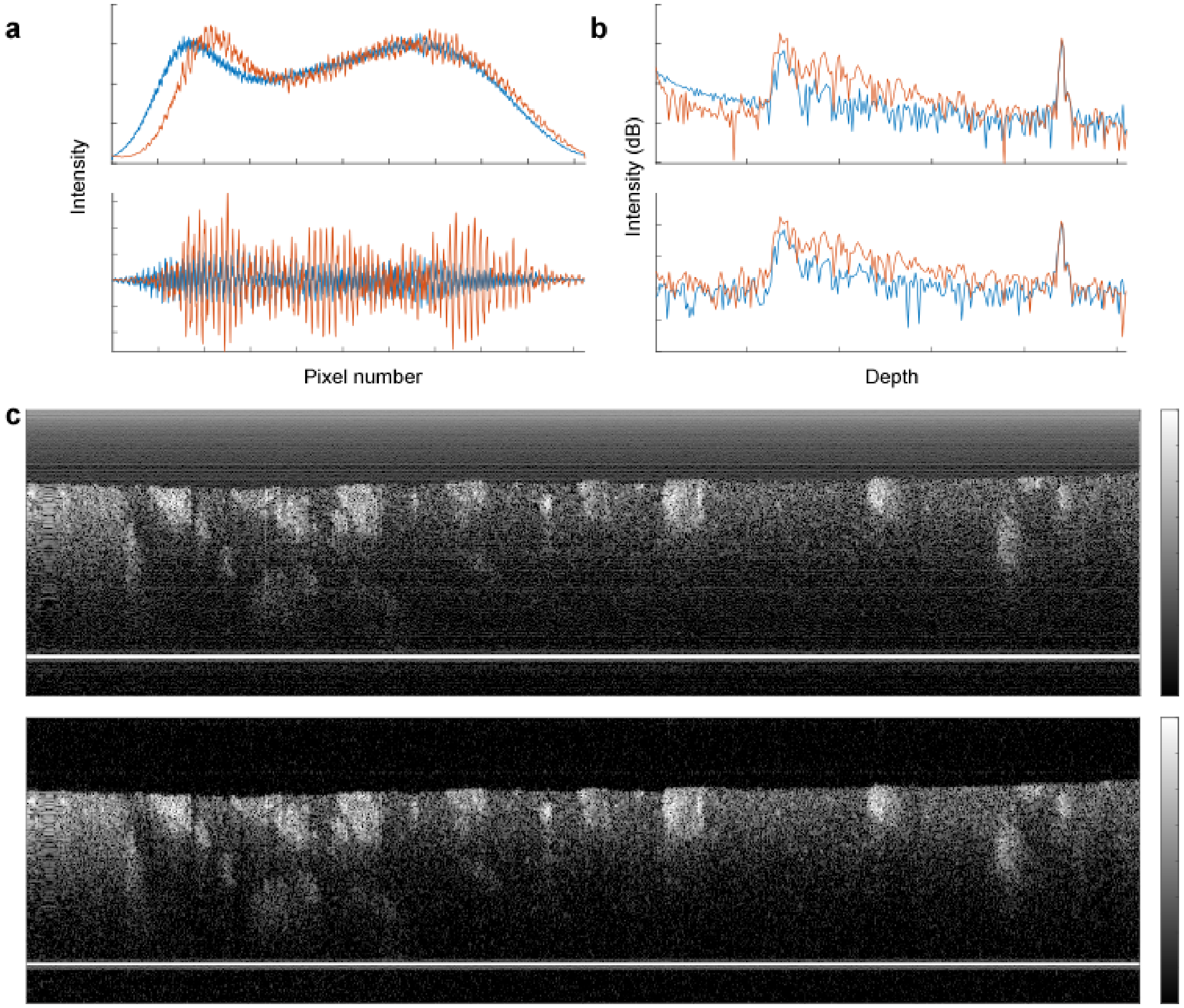


**Supplemental Figure 2: Background Subtraction. a.** Raw spectral data amplitude for a given camera pixel of one A-line, blue representing channel 2 and orange channel 1. **b.** The amplitude of the complex depth profile (the pre-processed A-line) against depth of the same A-line. **c.** Depth profiles in dB scale without and with background subtraction.

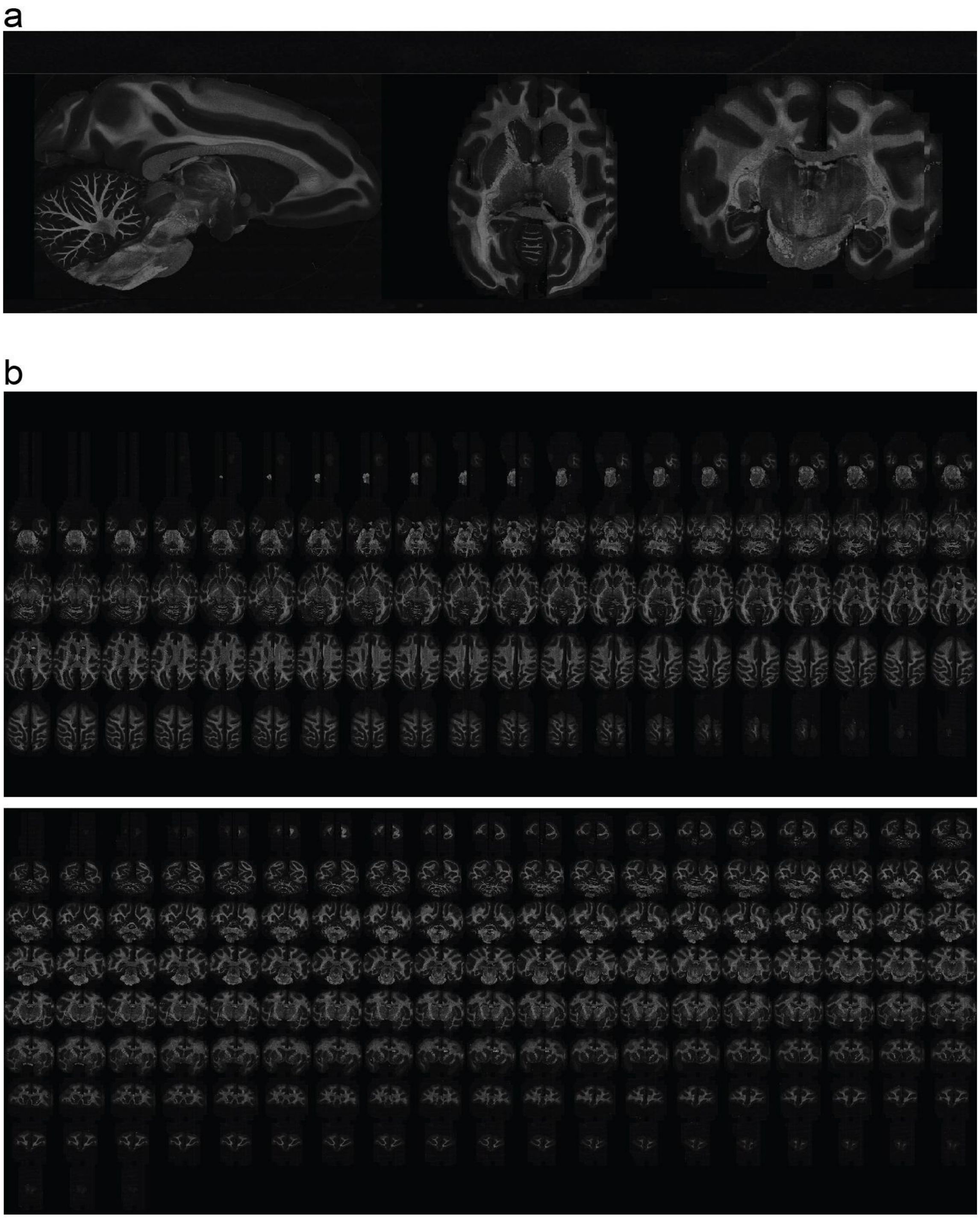


**Supplemental Fig. 3: Alignment across sections produces 3D PS-OCT volumes in monkey V.** After imaging in the sagittal plane, axial and coronal views show good between-slice alignment.

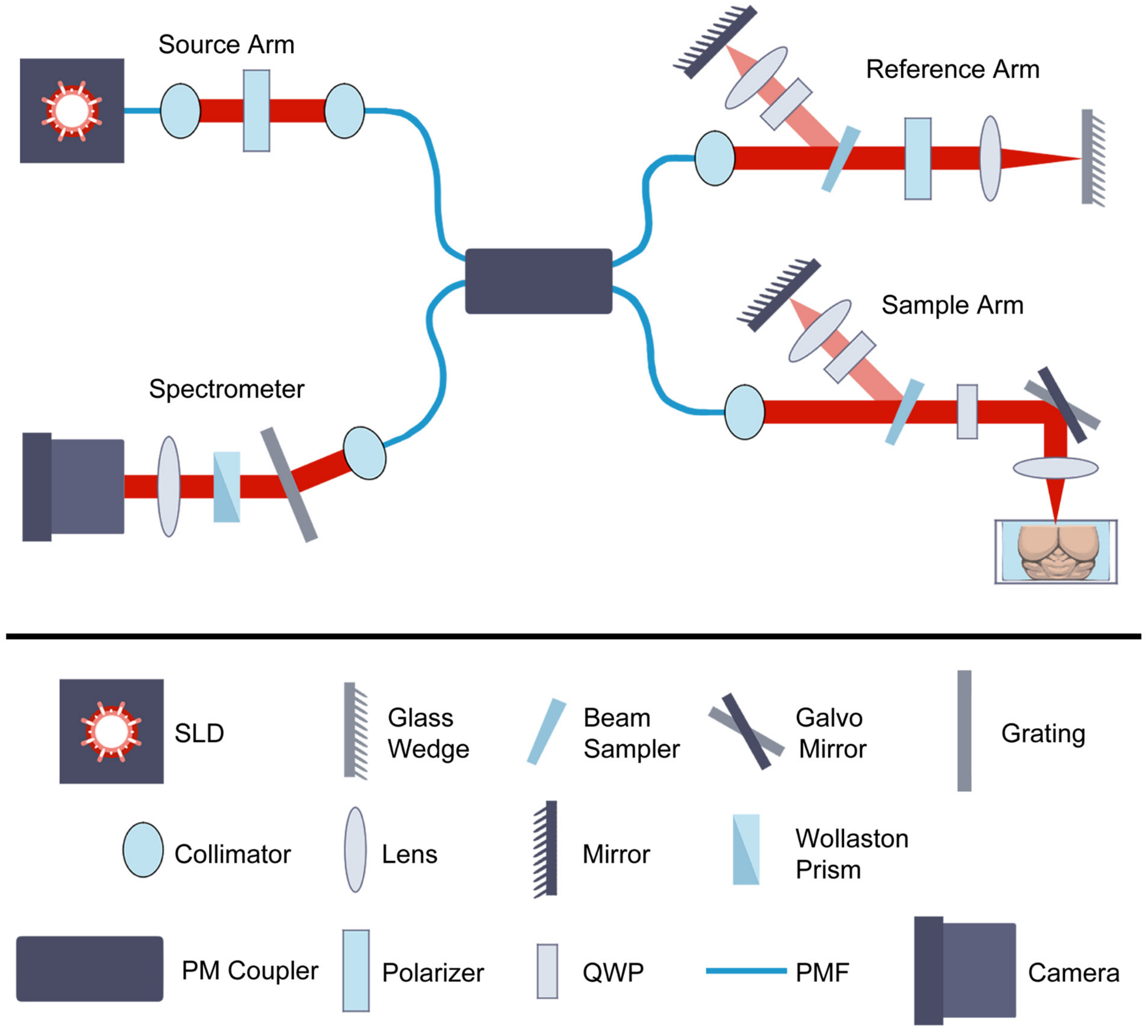


**Supplemental Figure 4: Optical system.** Detailed optical components of the interferometer. SLD: superluminescent diode, PM Coupler: polarization maintaining coupler, PMF: polarization maintaining fiber, QWP: quarter wave plate.
